# Fluid shear stress modulates endocytic pathways and junctional targeting of tumor-derived extracellular vesicles in endothelial cells

**DOI:** 10.64898/2026.05.01.721946

**Authors:** Nicolas Jones Villarinho, Bong Hwan Sung, Ana Sayuri Yamagata, Ramon Handerson Gomes Teles, Luiz Eduardo da Silva, André Zelanis, Murilo de Souza Salardani, Gisela Ramos Terçarioli, Vitória Samartin Botezelli, Julia de Moura Bernardi, Mario Costa Cruz, Ruy Gastaldoni Jaeger, Alissa Weaver, Vanessa Morais Freitas

## Abstract

Breast cancer is the most common malignancy in women, with triple-negative breast cancer (TNBC) representing the most aggressive subtype and carrying a poor metastatic prognosis. Metastasis requires tumor cells to cross the endothelial barrier, a process facilitated by tumor-derived extracellular vesicles (EVs), which can disrupt vascular integrity. Fluid shear stress (FSS), generated by blood flow, shapes endothelial physiology and may influence EV uptake, yet the mechanisms underlying TNBC-derived small EV (sEV) internalization remain unclear. Here, we investigated TNBC sEV–endothelial interactions using combined in silico and in vitro approaches. Human umbilical vein endothelial cells (HUVECs) were cultured under static or FSS conditions (20 dyn/cm²), followed by proteomic profiling and protein–protein interaction analyses with sEV proteomes. Uptake assays employed pharmacological inhibition (Dynasore, MβCD, Pitstop2), Caveolin-1 (CAV-1) and Clathrin Heavy Chain (CLHC), siRNA-mediated knockdown, and junctional interaction analyses via confocal microscopy and co-immunoprecipitation. FSS downregulated proliferation- and angiogenesis-associated proteins while upregulating adhesion and cytoskeletal regulators assessed by proteomics. Network analysis identified clathrin- and caveolin-mediated endocytosis (CME and CavME), integrins, and early endosomes as central mediators of sEV uptake. Functionally, uptake was reduced by Pitstop2, MβCD, and CAV-1/CLHC knockdown under static conditions, but silencing paradoxically enhanced uptake under FSS, suggesting compensatory flow-dependent pathways. Notably, under FSS, sEVs accumulated at endothelial junctions, colocalizing with VE-CAD and associating with CLDN5, indicating a potential disruption mechanism of adherens and tight junctions and consequent endothelial permeability. These findings identify CME and CavME as key uptake routes while underscoring FSS as a critical determinant of endothelial–tumor EV interactions. By revealing junctional targeting of sEVs, this work provides new mechanistic insight into vascular remodeling during metastasis and highlights EV pathways as potential therapeutic targets in TNBC.

**Graphical Abstract:** 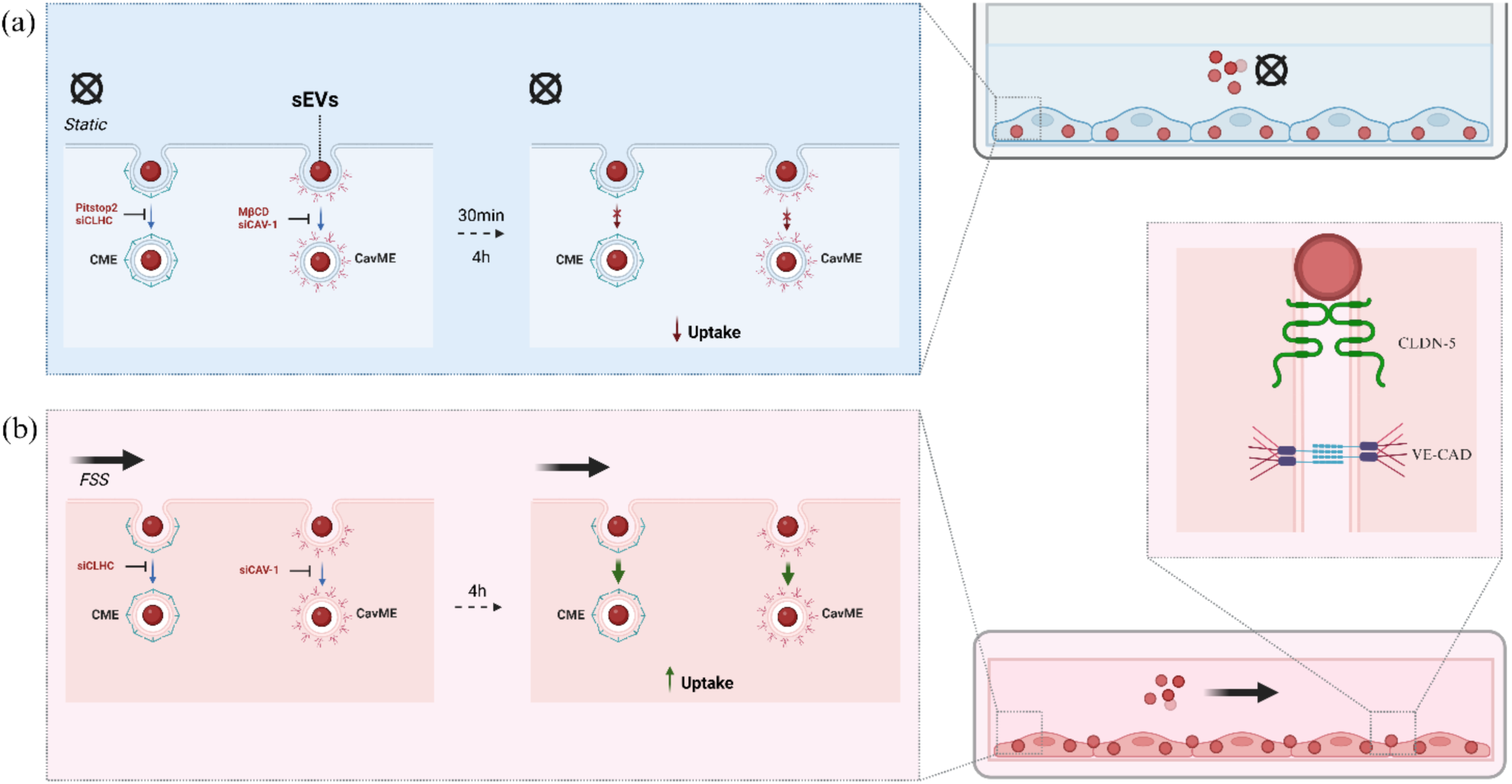

Uptake and localization of sEVs on HUVEC under (a) static and (b) fluid shear-stress conditions. sEVs: Small Extracellular Vesicles. CME: Clathrin-mediated Endocytosis. CavME: Caveolin-mediated Endocytosis. CLDN5: Claudin-5. VE-CAD: Vascular Endothelial Cadherin. FSS: Fluid shear-stress.

## 1. Introduction

Breast cancer is the most common type of cancer to affect women worldwide, being responsible for 2.0 million new cases (13.2% of all cancers, both sexes) and 1.2 million deaths (18.3% of all cancers, both sexes) in 2022 ^1^. Despite advances in screening and treatment, survival outcomes remain heterogeneous and are strongly influenced by tumor subtype and stage at diagnosis. Breast cancer is a highly heterogeneous disease driven by genetic, epigenetic, and environmental factors. Histopathological and molecular classifications provide the framework for prognosis and therapy. Based on hormone receptors (Estrogen and Progesterone) and HER2 expression, tumors are subdivided into luminal A, luminal B, HER2-enriched, and triple-negative breast cancer (TNBC) ^2,3^. Among them, TNBC accounts for 10-15% of cases and is characterized by the low expression of the hormone receptors and HER2. TNBC displays aggressive clinical behavior, high rates of relapses, and a strong propensity for visceral and brain metastases, with limited therapeutic options beyond chemotherapy and, more recently, immune checkpoint inhibitors and PARP inhibitors ^4,5^.

Metastasis is the leading cause of death in TNBC patients. This multistep process involves tumor cell dissemination from the primary site, intravasation, survival in the circulation, adhesion to the vascular endothelium, extravasation, and colonization of distant organs ^6,7^. Endothelial cells (ECs) are central players in these events, regulating vascular permeability, mediating tumor cell adhesion and transendothelial migration, and are themselves modulated by tumor-derived factors, such as extracellular vesicles (EVs) ^8,9^. ECs maintain vascular homeostasis through junctional complexes that regulate the barrier integrity. Adherens junctions (AJs), primarily mediated by VE-cadherin (VE-CAD), provide structural stability and control permeability, while tight junctions (TJs), composed largely of claudins, reinforce the paracellular barrier and preserve endothelial polarity ^10^. However, during the progress of tumorigenesis, ECs may undergo structural and functional alterations, leading to angiogenesis, increased leakiness, and facilitating tumor cell dissemination ^11,12^.

Recent evidence highlights the role of tumor-derived EVs in the metastatic cascade. These lipid bilayer–enclosed particles carry proteins, nucleic acids, and metabolites, and are key mediators of intercellular communication^13^. According to the 2023 Minimal Information for Studies of EVs (MISEVs 2023), EVs can be categorized according to their biogenesis (e.g., exosomes originate from the fusion of multivesicular bodies with the plasma membrane) or by size, being referred to as small EVs (sEVs, <200 nm in diameter) or large EVs (>200 nm). In cancer, EVs promote tumor progression by enhancing angiogenesis, immune evasion, and vascular permeability, as well as by preparing pre-metastatic niches ^14–16^. Importantly, EV uptake by ECs has been shown to occur through multiple endocytic pathways, including Caveolin and Clathrin–mediated endocytosis (CavME and CME), and may directly impact endothelial junctional integrity ^17–21^. Despite growing evidence, the molecular mechanisms underlying how tumor-derived EVs regulate EC biology and facilitate TNBC metastasis remain incompletely understood.

ECs, by lining the lumen of blood and lymphatic vessels, are continuously exposed to mechanical stresses that vary according to vessel type and caliber ^22^. These forces can be broadly classified as contact-derived stresses, arising from vessel topography, curvature, and stiffness, or flow-derived stresses, including tension, pressure, and fluid shear stress (FSS) ^23^. FSS represents a frictional force exerted by blood flow on the endothelial surface ^24^ and its magnitude differs across the vasculature, ranging from 1–22 dyn/cm² in the aorta, 10–70 dyn/cm² in arteries, 1–6 dyn/cm² in veins, and 3–95 dyn/cm² in capillaries ^25–28^. ECs sense and transduce these hemodynamic cues through mechanotransduction pathways involving the plasma membrane, adhesion sites, and the cytoskeleton ^29,30^. In vitro, ECs cultured under static conditions display an epithelioid morphology, whereas exposure to FSS induces elongation and cytoskeletal alignment in the direction of flow ^31^. Additional mechanosensors include ion channels (e.g., Ca²⁺ channels), cell–cell junctional complexes such as VE-cadherin and PECAM-1, primary cilia, caveolae, and cytoskeleton-associated structures, which together form an interconnected network linking external biomechanical forces to intracellular responses ^32^.

These mechanotransductive processes are crucial for endothelial functions such as migration and adhesion and have been implicated in cancer progression. From a biomechanical perspective, FSS influences metastatic dissemination by regulating the adhesion of circulating tumor cells (CTCs) to the vascular endothelium. In brain metastases, low shear stress levels typical of capillaries enhance CTC retention, adhesion, and extravasation, whereas higher shear forces, such as those in arteries, reduce adhesion. Conversely, extremely low shear conditions favor CTC adhesion but impair their extravasation ^33,34^. Moreover, FSS can activate signaling pathways, including NF-κB, PI3K/AKT, and c-Jun, and induce the expression of cytokines, integrins, and mechanosensitive molecules that collectively promote invasion and metastasis. However, it is still unclear how FSS affects the uptake and intracellular localization of tumor-derived EVs in ECs.

In this study, we investigated the interaction and uptake of sEVs by ECs using a combination of *in silico* and *in vitro* approaches. We first compared the proteomic profiles of HUVECs cultured under static conditions or exposed to FSS. In parallel, we characterized the proteomic composition of sEVs derived from the TNBC cell line MDA-MB-231. Protein–protein interaction analyses were then performed to predict potential interactions between HUVECs (with or without FSS exposure) and sEVs, focusing on endocytic pathways. To functionally validate these predictions, we modulated clathrin- and caveolin-mediated endocytosis in HUVECs under static and FSS conditions and assessed their impact on sEVs uptake. Notably, sEVs preferentially accumulated at cell–cell junctions under FSS, prompting further investigation into the junctional proteins mediating this interaction. These findings provide critical insight into the mechanobiological regulation of sEVs uptake by ECs, with implications for understanding the early steps of TNBC metastasis and identifying potential therapeutic targets.

## 2. Results

### 2.1. Comparison of the Proteomic Profile of HUVEC under Static and FSS Conditions

To explore potential endocytic pathways for sEVs uptake, we performed proteomic profiling of HUVECs cultured under static or FSS conditions. Cells were maintained in 4% dextran to mimic blood viscosity, with FSS exposure at 20 dyn/cm² via orbital shaking for 96 h (Figure 1A), while static controls were cultured without agitation.

**Figure 1:**
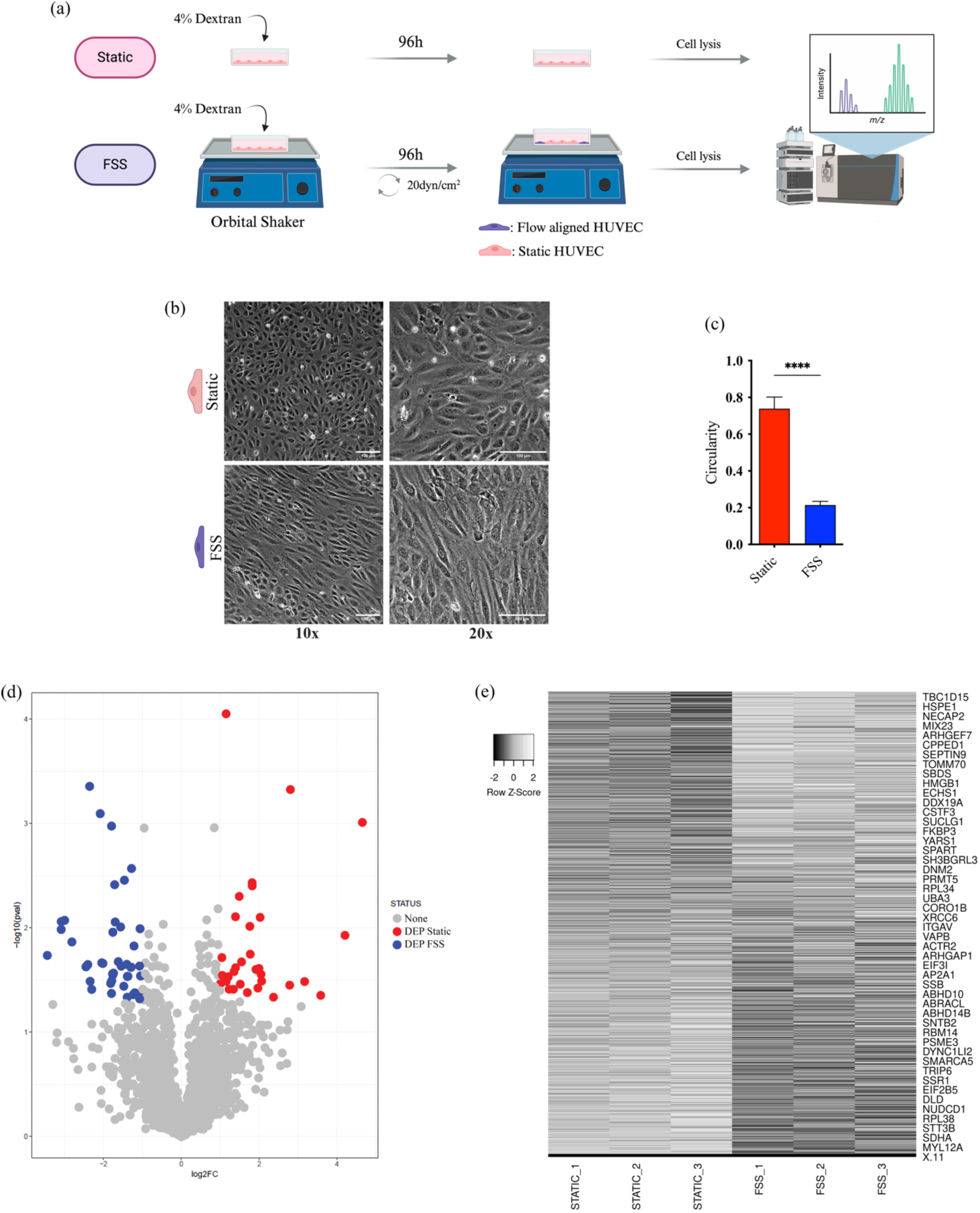
FSS induces morphological and proteomic changes in HUVECs. (a) Schematic representation of HUVEC culture under static or FSS conditions through the orbital shaker method, followed by proteomic analysis using LC–MS/MS. (b) Phase-contrast micrographs of HUVECs aligned (FSS) or not aligned (Static) under FSS. (c) Cell circularity assessed by the width-to-length ratio of individual cells. (d) Volcano plot showing proteins differentially expressed between conditions. (e) Heatmap showing differential protein expression between STATIC and FSS conditions. Expression values are presented as row Z-scores. FC: fold-change; Pval: p-value; DEP: differentially expressed proteins; None: proteins not altered by FSS. Data are from three independent experiments and are presented as mean ± SEM. Statistical analysis was performed using Student’s *t*-test. ****: p ≤ 0.001.

After 96 h, static HUVECs were cuboidal with high circularity (0.74 ± 0.34 A.U.), whereas FSS-exposed cells were elongated, aligned with flow, and showed reduced circularity (0.21 ± 0.11) (Figures 1B-C). Proteomic analysis identified 32 DEPs in static cells and 43 in FSS-exposed cells (Figure 1D, full lists in Supplementary Tables 1-2). The heatmap shows distinct gene expression profiles under static and FSS conditions, with clear segregation between groups indicating condition-specific transcriptional responses (Figure 1E). Static HUVECs exhibited upregulation of endocytosis-related proteins, including ANXA7, ANXA11, and MYO6, whereas cells exposed to FSS displayed enrichment in pathways associated with lysosomal transport (LAMP1), cell cycle arrest (HMGA2, RSF1, CEP164), anti-proliferative activity (IGFBP4), and metabolic regulation (C11orf54, CPT2) (Supplementary Tables 1-2).

These findings suggest that static HUVECs are more prone to particle internalization, whereas FSS-exposed cells, under shear mimicking large vessels, reduce proliferation and alter metabolism. GO enrichment analysis of the top 20 biological processes revealed that static cells favor angiogenesis, proliferation, adhesion, migration, and endocytosis, whereas FSS-exposed cells emphasize adhesion, cytoskeletal organization, polarity, and growth response, highlighting distinct phenotypes adapted to flow conditions (Figures 2A–B).

**Figure 2:**
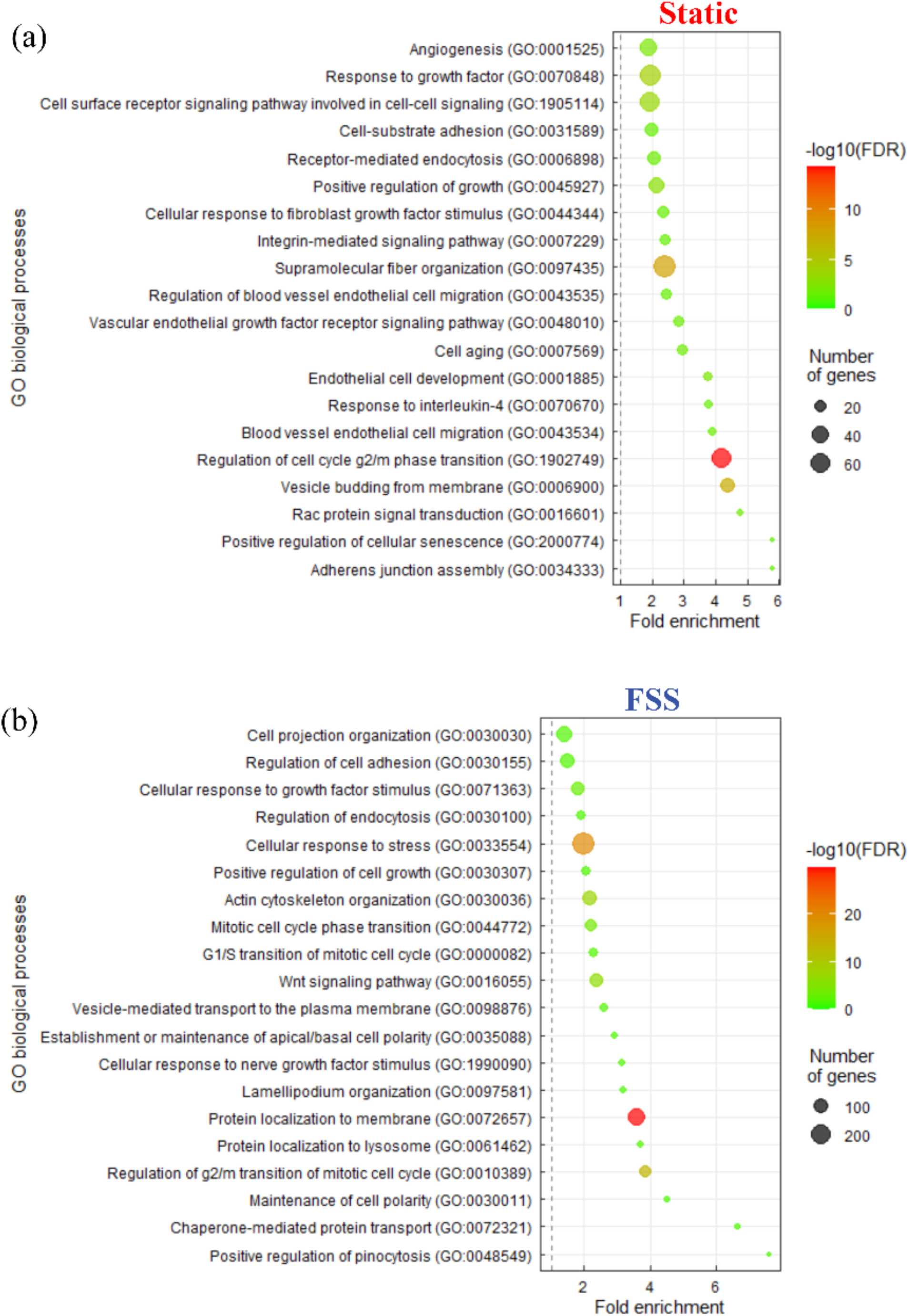
FSS alters the biological processes associated with DEPs in HUVECs. GO enrichment analysis of HUVECs under (a) static and (b) FS conditions. GO: Gene Ontology. FDR: False Discovery Rate.

### 2.2. Characterization of MDA-MB-231 CD63-mScarlet sEVs

Prior to the in silico analysis, EVs were isolated using Cushioned Density Gradient Ultracentrifugation, in which sEVs migrate according to their buoyant density within the gradient and are typically recovered in fractions 6 and 7. (Figure 3A) ^35^. The sEVs were validated by NTA, TEM, Western blotting, and confocal microscopy. To enable vesicle tracking in endocytosis assays, MDA-MB-231 cells were engineered to stably express CD63 fused to the fluorescent protein mScarlet, generating the MDA-MB-231 CD63-mScarlet cell line. Success of transfection was verified by flow cytometry, which revealed a cellular profile comparable to the parental line and a marked increase in fluorescence intensity, confirming efficient gene delivery (Supplementary Figure 1). This approach confined fluorescence to EVs, avoiding nonspecific labeling associated with membrane dyes and improving assay reliability.

**Figure 3:**
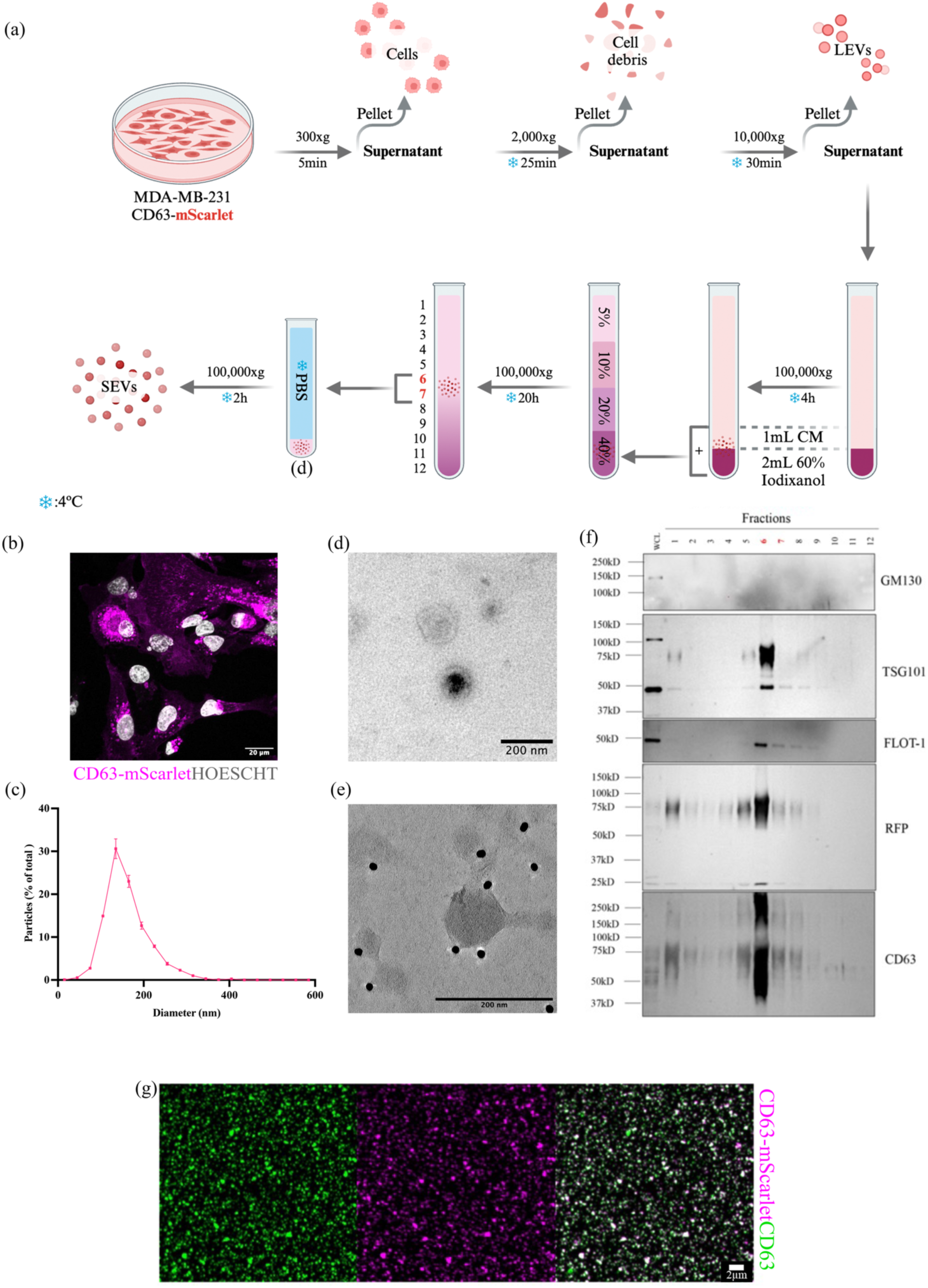
Characterization of sEVs derived from the MDA-MB-231 cell line expressing the CD63-mScarlet fusion protein. (a) Schematic of EV isolation using Cushioned Density Gradient Ultracentrifugation. Illustration created with BioRender.com. Red numbers indicate localization of sEVs within the gradient. (b) Transfection of MDA-MB-231 cells with the CD63-mScarlet plasmid was assessed by confocal microscopy. (c) NTA of particles from fractions 6 and 7 of the gradient. (d) Transmission electron microscopy of sEVs and € immunogold labeling of the sEVs marker CD63. (f) Western blotting of whole-cell lysate (WCL) and iodixanol discontinuous gradient fractions, probed with antibodies against EV markers (TSG101, FLOT-1, CD63), the fluorescent marker (RFP), and the EV-negative marker GM130. (g) Adhesion assay of sEVs on glass coverslips, showing fluorescence from CD63-mScarlet and immunolabeling with anti-CD63.

In the transfected cell line, cytoplasmic expression of CD63-mScarlet was observed, reflecting its localization to multivesicular bodies (Figure 3B). Characterization of sEVs derived from this cell line followed the MISEV 2023 guidelines and revealed that the combined fractions 6 and 7 contained particles with a mean diameter of 161.1 ± 51.27 nm (Figure 3C), exhibiting the characteristic cup-shaped morphology (Figure 3D) and positive labeling for CD63 under TEM (Figure 3E). An enrichment of EV markers (TSG101, FLOT-1, and CD63) was observed in fractions 6 and 7 of the continuous gradient, together with mScarlet (Figure 3F). GM130 was only detected in the WCL, confirming the absence of contaminating cellular debris in the sEVs samples (Figure 3F). Moreover, mScarlet fluorescence colocalized with CD63 immunostaining, confirming successful fusion of CD63 with mScarlet (Figure 3G).

This characterization served as the foundation for the subsequent *in silico* and, later, *in vitro* endocytosis assays. For the following experiments, fractions 6 and 7 were pooled and quantified.

### 2.3. In silico characterization of MDA-MB-231 CD63-mScarlet sEVs

Given the alterations observed in HUVEC under different levels of FSS, we next examined their interaction with MDA-MB-231 CD63-mScarlet–derived sEVs. To validate the proteomic content, we compared our sEVs protein datasets with Vesiclepedia (Figure 4A, Supplementary Table 3). Among sEVs, 87.5% matched previously reported entries.

**Figure 4:**
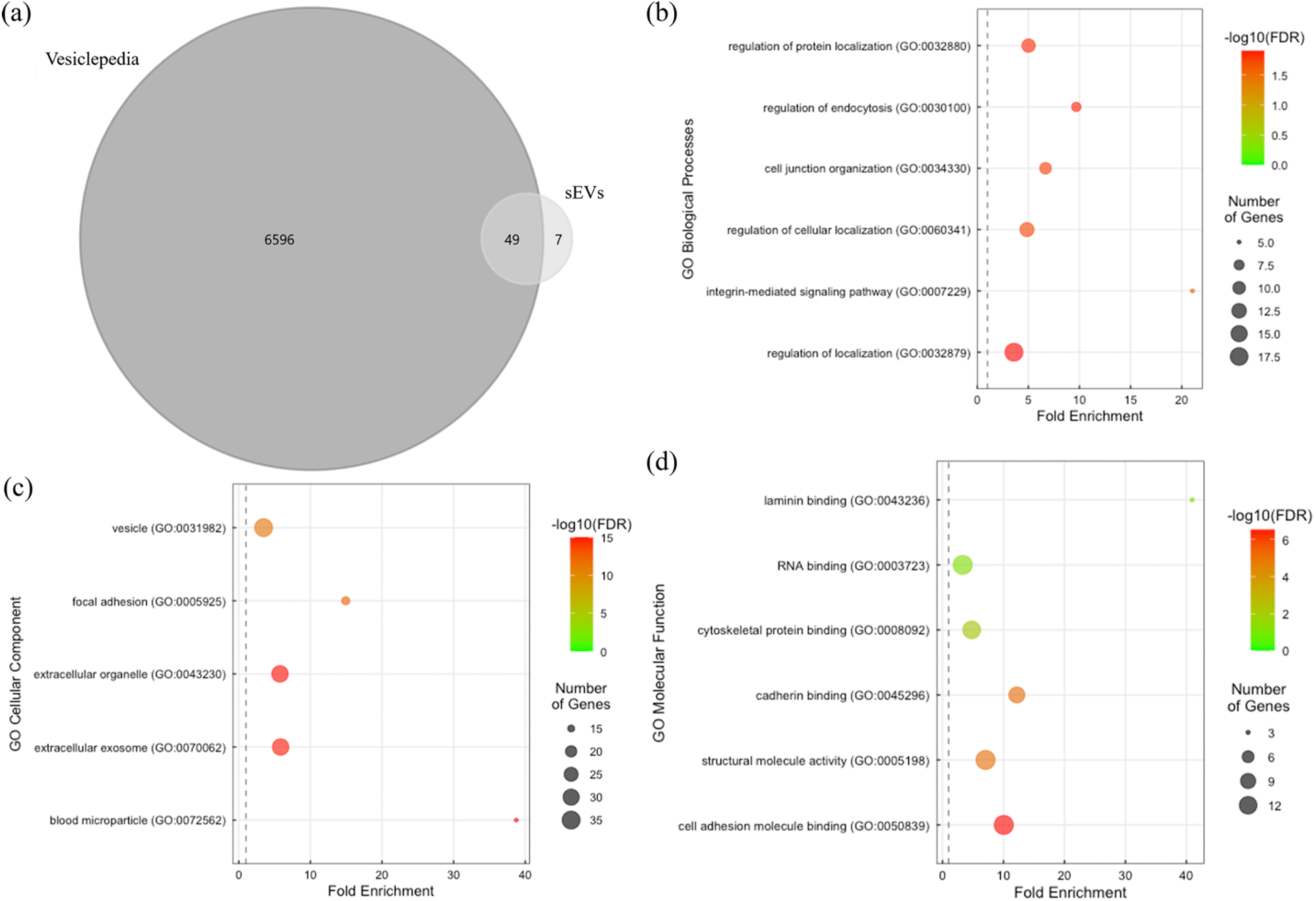
GO enrichment analysis of sEVs derived from the MDA-MB-231 cell line. (a) Proteomic profile of sEVs compared with all proteins reported in the Vesiclepedia database. Gene Ontology (GO) enrichment analysis of sEVs highlighting terms related to (b) biological processes, (c) cellular components, and (d) molecular functions. GO: Gene Ontology. FDR: False Discovery Rate.

We next performed GO enrichment analysis of sEVs proteins. These proteins were mainly associated with biological processes such as cellular localization, endocytosis, cell junction organization, and integrin-mediated signaling (Figure 4B). In terms of cellular components, they localized to extracellular vesicles, focal adhesions, and blood microparticles (Figure 4C). Molecular function analysis revealed associations with laminin binding, RNA binding, cytoskeletal interactions, cadherin binding, cell adhesion molecule binding, and structural components (Figure 4D). These findings are consistent with the expected profile of sEVs proteins and highlight the diverse functional roles of these particles.

### 2.4. Protein-Protein-Interaction (PPI) analysis between sEVs and HUVECs in the context of endocytosis

Based on the proteomic profiles of HUVECs under static and FSS conditions, together with MDA-MB-231 sEVs proteomics, we constructed an interactome to explore potential mechanisms of MDA-MB-231 sEVs uptake. HUVEC proteins common to both static and FSS conditions were combined with differentially expressed proteins (DEPs) from each group. GO enrichment analysis was then applied, filtering only proteins annotated to Endocytosis (GO:0006897). Both conditions showed similar profiles, except for MYO6, which was among the top 10 expressed proteins under static conditions.

PPI analysis revealed three proteins shared by HUVECs (static and FSS) and MDA-MB-231 sEVs (ITGB1, ACTB, ACTG1), as well as sEVs interactions with clathrin-mediated endocytosis (CME, CLTA, CLTC) and caveolin-mediated endocytosis (CavME; CAV1) (Figure 5A). Notably, CLTA interacted with MYO6 in static HUVECs, while MYO6 also interacted with the MDA-MB-231 sEVs protein MYL6, suggesting a potential role for MYL6 in TNBC cancer cells sEVs uptake (Figure 5A). In addition, MDA-MB-231 sEVs integrins (ITGB1, ITGA3, ITGA6) were linked to CavME, suggesting an uptake route. Network centrality analysis further highlighted CME proteins as the most connected nodes (Figure 5B–C). Together, these findings suggest that both CME and CavME pathways contribute to MDA-MB-231 sEVs internalization in HUVECs under static and FSS conditions.

**Figure 5:**
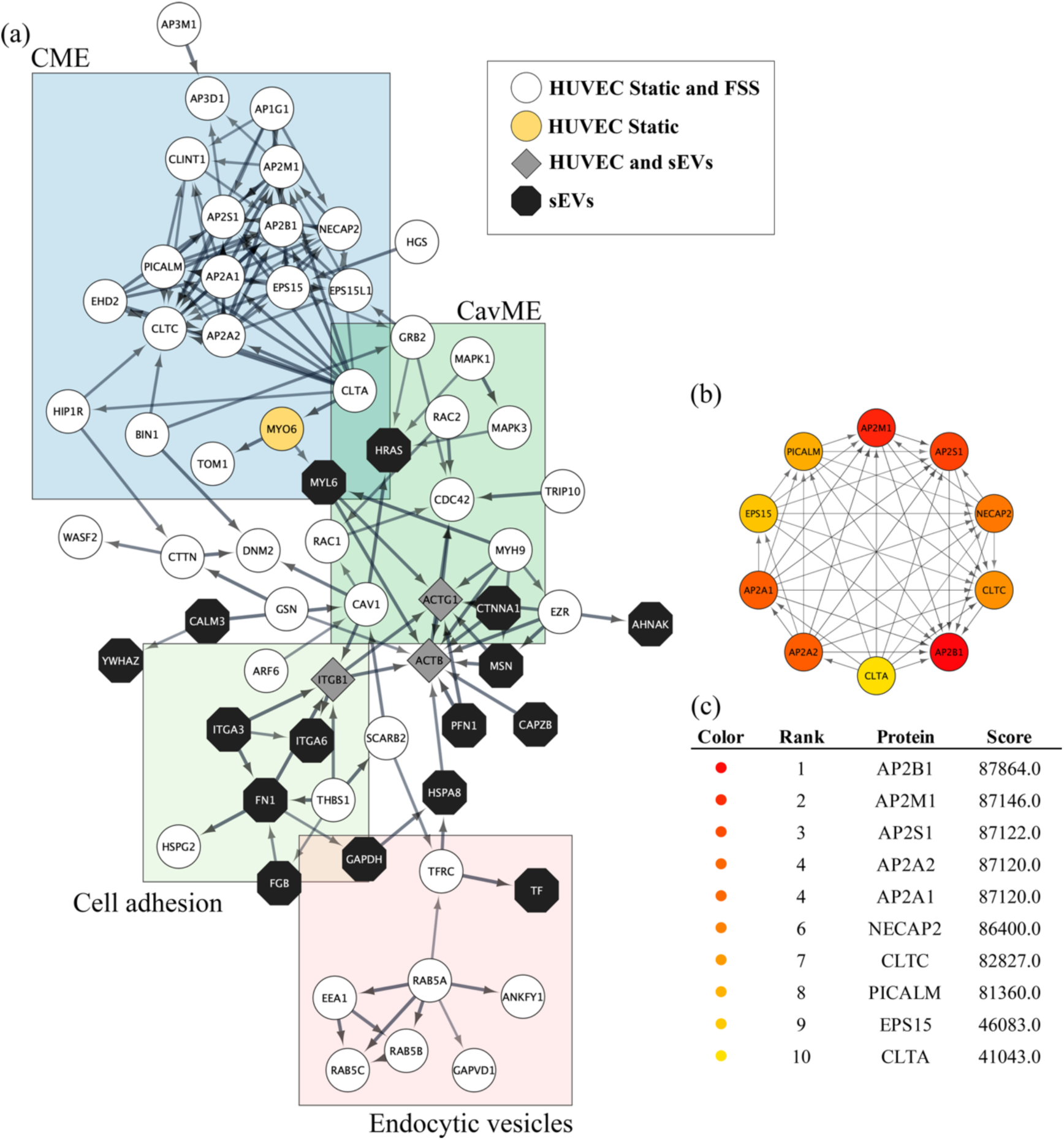
Interactome highlights CME- and CavME-mediated endocytosis in the interaction between MDA-MB-231 sEVs and HUVECs under static and FSS conditions. (a) Network representation of endocytosis-related proteins in HUVECs under static and FSS conditions (white circles), proteins exclusive to static HUVECs (yellow circles), shared HUVEC-sEV proteins (gray diamonds), and sEV-exclusive proteins (black octagons). Arrows indicate interaction direction (upstream x downstream), arrows thickness is related to PPI score (0.4 to 1.0) and colored boxes highlight nodes linked to specific cellular mechanisms. (b) Central node interactions ranked by Maximal Clique Centrality (MCC). (c) Top 10 hub nodes of the interactome.

### 2.5. MβCD and Pitstop2 inhibit MDA-MB-231 sEVs uptake in static HUVEC

After validating CD63-mScarlet-labeled MDA-MB-231 sEVs and selecting pharmacological inhibitors for endocytosis, we first tested their effects on MDA-MB-231 sEVs uptake in static HUVEC culture. The static condition was chosen because acid wash and inhibitors disrupted cell–cell adhesion, making us test FSS with transient cell transfection, as further described. Cells were then pretreated with Dynasore (100μM), MβCD (2.5μg/mL), or Pitstop2 (30μM).

MβCD significantly reduced MDA-MB-231 sEVs uptake at both 30 min (53%) and 4 h (48%), while Pitstop2 reduced uptake only at 30 min (83%) (Figure 6A-B), supporting roles for both CME and CavME in sEVs internalization. Interestingly, prolonged Pitstop2 treatment increased sEVs accumulation after 4 h, coinciding with HUVEC morphological changes, such as loss of stress fibers and punctate cytoskeletal staining, suggesting cell death-related retention of MDA-MB-231 sEVs.

**Figure 6.**
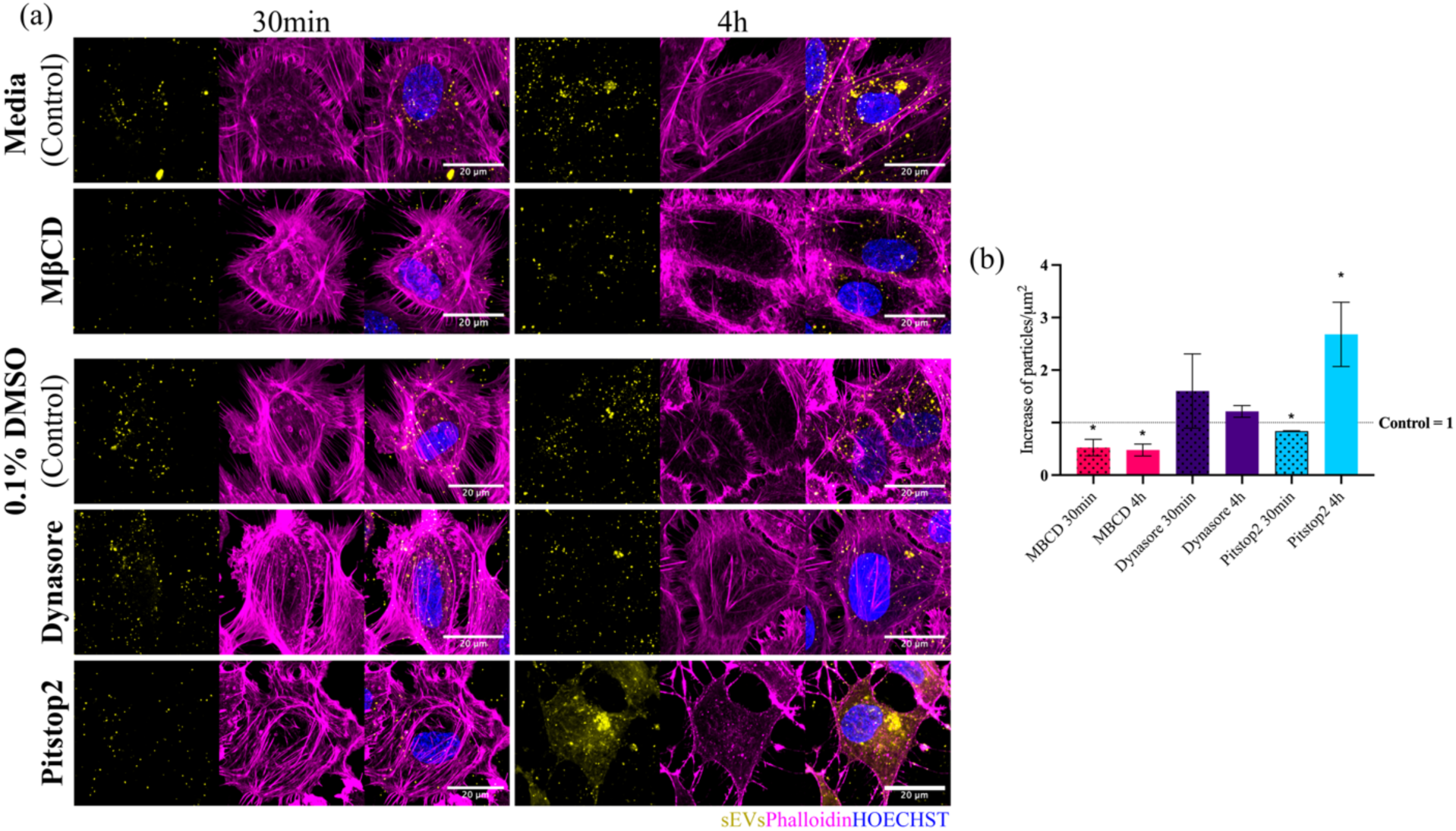
MβCD and Pitstop2 inhibit MDA-MB-231 sEV uptake in static HUVECs. (a) Effect of endocytosis inhibitors MβCD, Dynasore, and Pitstop2 after 30 min and 4 h of incubation with sEVs. Vehicle served as control. (b) Quantification of sEV particles per area relative to control. Data represent mean ± SEM from three independent experiments. Statistical analysis by one-way ANOVA. *p ≤ 0.05. Scale bar: 20μm.

### 2.6. Knockdown of CAV-1 and CLHC reduces sEVs uptake in static HUVEC but not under FSS

Given that both CME and CavME appeared to influence MDA-MB-231 sEVs uptake by endothelial cells, we next examined how their inhibition would affect endocytosis in HUVEC under FSS, a condition more representative of the in vivo environment. To avoid the cytotoxic effects of pharmacological inhibitors, transient knockdown of CAV-1 and CLHC was performed using siRNA. Forty-eight hours post-transfection, CAV-1 expression was reduced by 60% and CLHC by 48% (Supplementary Figure 6A–C). Notably, CLHC knockdown also decreased CAV-1 levels by 35% (Supplementary Figure 6C).

We investigated how FSS affects MDA-MB-231 sEVs uptake in HUVEC transiently transfected with siRNA targeting CAV-1 or CLHC. HUVEC were aligned under laminar flow (20 dyn/cm², 24h) and exposed to sEVs under reduced FSS (8.5 dyn/cm²) in µ-slide y-shaped channels simulating High FSS (8.5 dyn/cm²) in the first straight line until the bifurcation (BIF), and from the convergence (CONV) along the second straight line, while the Low FSS (4.25 dyn/cm²) was achieved in the curved zone indicated by the parallels central lines (Figure 7B and 8B). FSS values were determined based on the findings of Mack et al. (2017), taking into account that CTC adhesion and extravasation typically occur within the microvasculature, where FSS is relatively low ^37,38^. Static conditions were analyzed in parallel.

**Figure 7.**
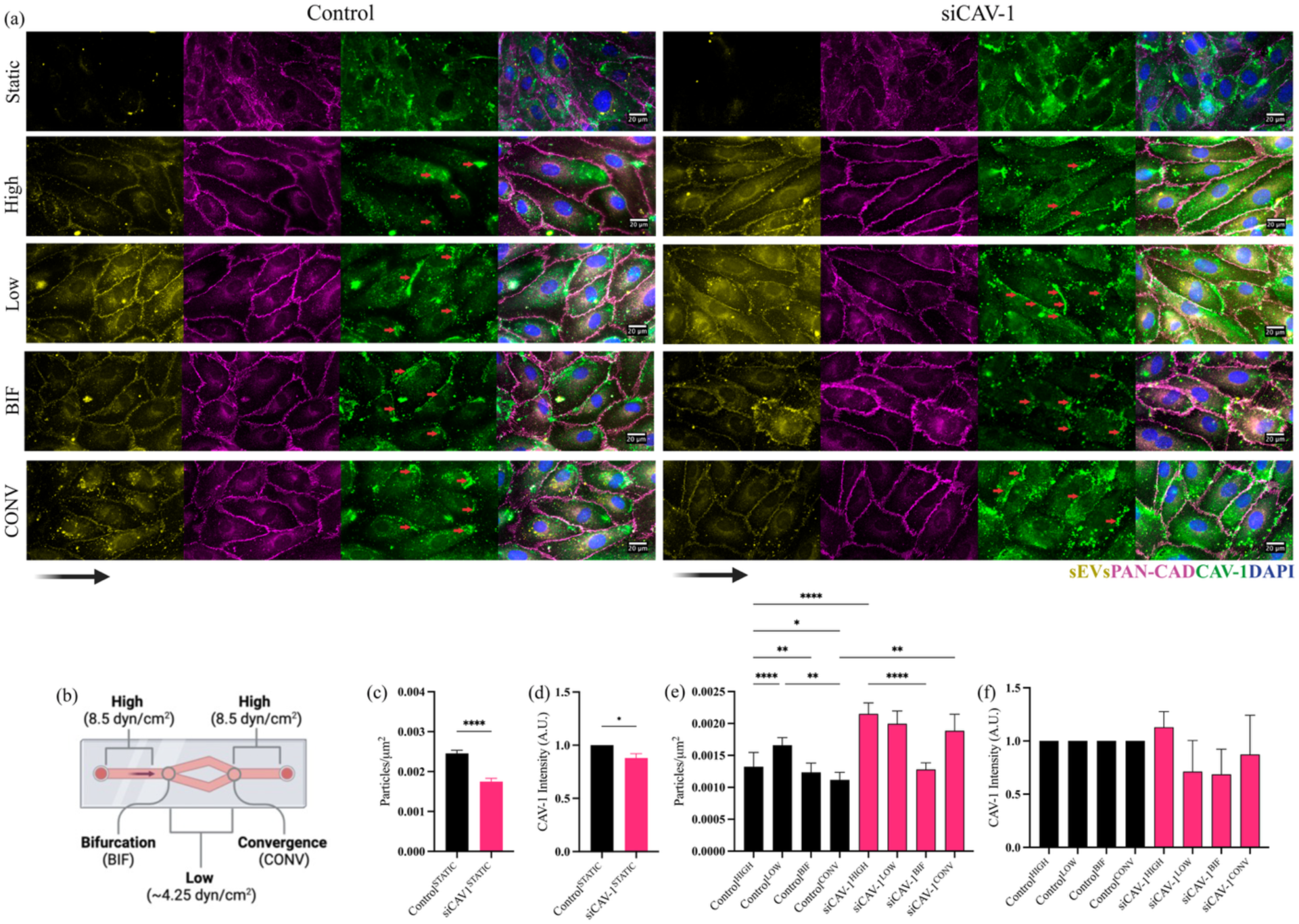
CAV-1 knockdown reduces MDA-MB-231 sEVs uptake under static conditions but increases it under FSS. (a) Effect of siRNA-mediated CAV-1 knockdown (siCAV-1) in HUVECs under static and FSS conditions assessed by fluorescence microscopy. White arrows indicate CAV-1 deposition downstream of laminar flow; the black arrow indicates the flow direction. (b) Schematic representation of the Y-shaped µ-slide showing the different regions analyzed. The arrow indicates the direction of laminar flow. Illustration created using BioRender.com based on Mack *et al.,* 2017. (c–d) Quantification of particle uptake per area (c) and CAV-1 protein levels (d) under static conditions. (e–f) Quantification of particle uptake per area (e) and CAV-1 protein levels (f) under FSS. Results represent three independent experiments. Statistical analysis was performed using Kruskal–Wallis test. *p ≤ 0.05, **p ≤ 0.01, ****p ≤ 0.0001. sEVs: Small Extracellular Vesicles; CAV-1: Caveolin-1; PAN-CAD: Pan-cadherin; CLHC knockdown decreased MDA-MB-231 sEVs uptake under static conditions, without significantly altering CLHC expression (Figures 8A, C, D). Under FSS, siCLHC increased MDA-MB-231 sEVs internalization in all regions, indicating a compensatory mechanism under flow (Figures 8A, C, E).

**Figure 8.**
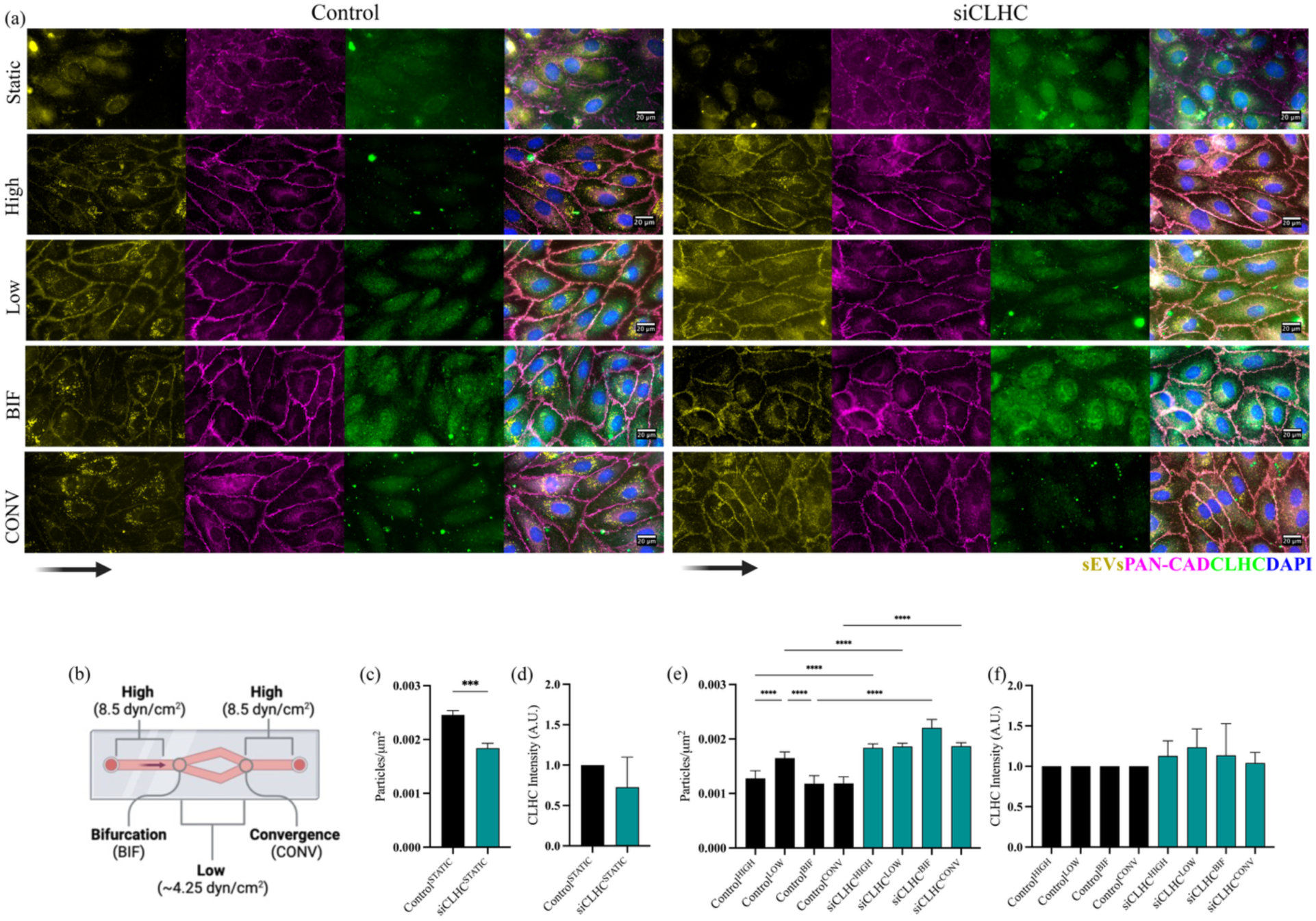
CLHC knockdown reduces MDA-MB-231 sEVs uptake under static conditions but increases it under FSS. (a) Effect of siRNA-mediated CLHC knockdown (siCLHC) in HUVECs under static and FSS conditions assessed by fluorescence microscopy. The black arrow indicates the flow direction. (b) Schematic representation of the Y-shaped µ-slide showing the different regions analyzed. The arrow indicates the direction of laminar flow. Illustration created using BioRender.com based on Mack *et al.,* 2017. (c–d) Quantification of particle uptake per area (b) and CLHC protein levels (c) under static conditions. (e–f) Quantification of particle uptake per area (e) and CLHC protein levels (f) under FSS. Results represent three independent experiments. Statistical analysis was performed using Kruskal–Wallis test. *p ≤ 0.05, **p ≤ 0.01, ****p ≤ 0.0001. sEVs: Small Extracellular Vesicles; CLCH: Clathrin Heavy Chain; PAN-CAD: Pan-cadherin; These results indicate that CAV-1 and CLHC govern sEVs uptake in static HUVEC, whereas under FSS, flow-mediated mechanisms may compensate for knockdown, increasing sEVs internalization. Notably, sEVs preferentially accumulate at cell-cell junctions only in HUVEC exposed to FSS, suggesting these sites may facilitate circulating tumor cell adhesion or contribute to enhanced vascular permeability.

Under FSS, HUVEC were elongated and Pan-cadherin localized at cell–cell junctions; static condition showed cuboidal cells with diffuse junctions (Figures 7A, 8A). CAV-1 knockdown reduced MDA-MB-231 sEVs uptake in static cells, correlating with decreased CAV-1 staining (Figures 7C, D). Under FSS, MDA-MB-231 sEVs accumulated at junctions in all groups, and CAV-1 localized downstream (Figure 7A, red arrows), indicating flow-dependent regulation. Uptake was highest in Low regions in controls; siCAV-1 increased MDA-MB-231 sEVs internalization under FSS, except at BIF, suggesting FSS compensates for knockdown (Figures 7E, F).

### 2.7. MDA-MB-231 sEVs colocalize with cell-to-cell junctions and interact with CLDN-5

Building on previous observations of sEVs accumulation at HUVEC cell-cell junctions under FSS, we validated these findings using confocal microscopy and Co-IP assays to assess interactions between MDA-MB-231-derived sEVs proteins and endothelial CLDN-5 under static and FSS conditions. sEVs were found to co-localize with the AJ protein VE-CAD mostly under FSS, validating our previous results (Figure 9A). Notably, mScarlet (detected with a dsRed2 antibody) co-precipitated with OJ protein CLDN-5 in FSS-exposed HUVEC (Figure 9B), suggesting that tumor-derived sEVs may directly target occludens junctions, revealing a potential mechanism of action.

**Figure 9.**
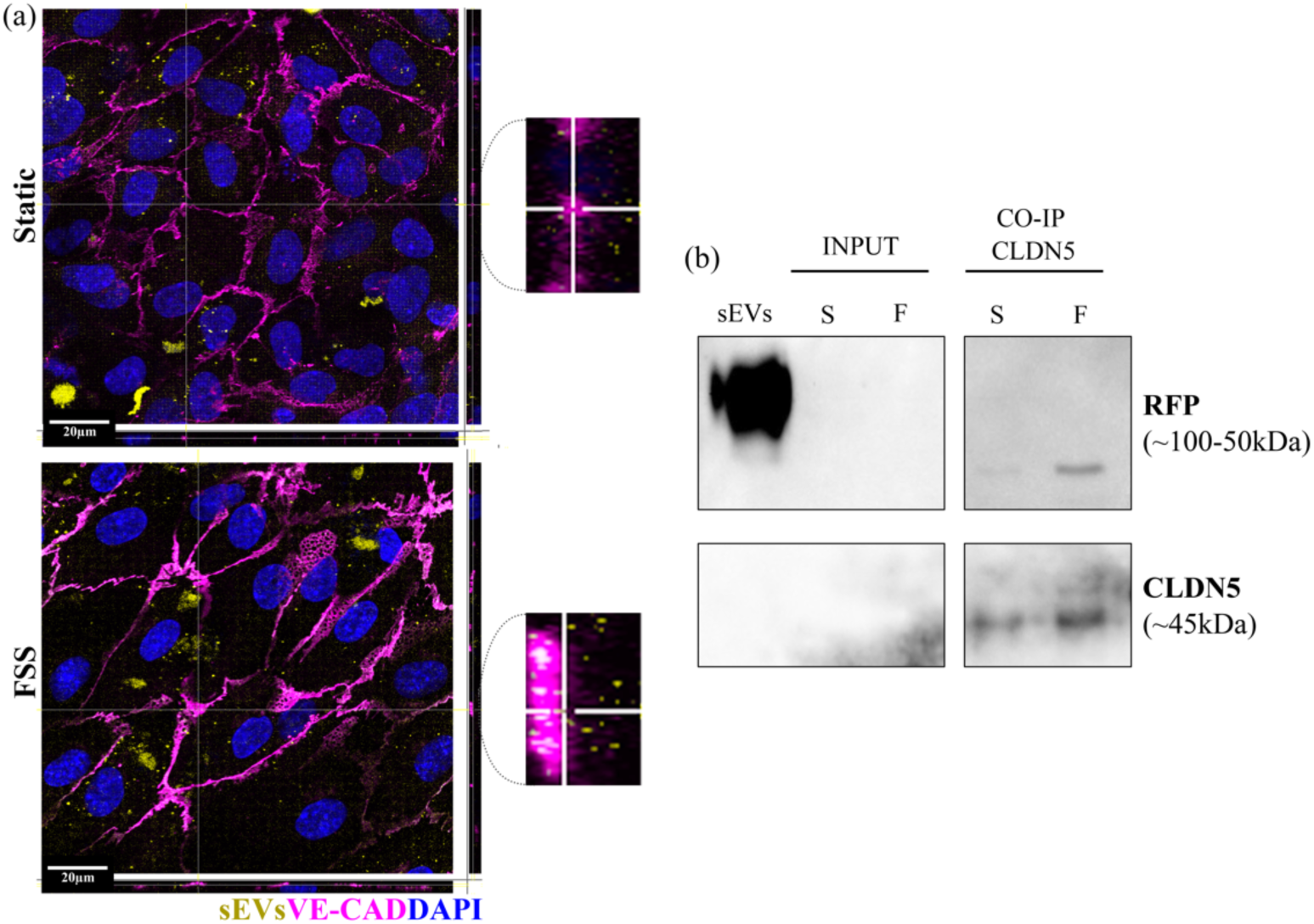
MDA-MB-231 sEVs colocalize with endothelial VE-CAD and interact with CLDN5 only under FSS. The panel shows HUVECs under FSS with immunostaining for VE-CAD, highlighting adherens junctions. YZ projections are shown to the right and XZ projections at the bottom of each image. (a) Field showing sEVs localization in HUVEC cells under static and FSS conditions. Images were acquired by confocal microscopy. Inserts highlights localization of sEVs on endothelial AJ. (b) Co-immunoprecipitation demonstrating that sEV-associated dsRed (mScarlet) interacts with endothelial CLDN5 under FSS. sEVs: Small vesicles, S: Static, F: FSS. Scale bar = 20µm

## 3. Discussion

In this study, we investigated the interaction between tumor-derived sEVs and ECs under static and FSS conditions, combining *in silico* and *in vitro* approaches to identify proteins involved in uptake and to clarify the endocytic pathways engaged. Proteomic and morphological analyses revealed that HUVECs adapt their phenotype to FSS by downregulating proteins associated with proliferation and angiogenesis while upregulating those related to cytoskeletal organization, adhesion, and structural maintenance. In silico protein-protein interaction analyses further suggested that CME and CavME, along with integrins and early endosomes, represent key routes for sEVs uptake.

Omics approaches are powerful tools to map cellular adaptations and guide mechanistic studies. In ECs, previous integrative proteomic and metabolomic studies have shown that low FSS suppresses lipid metabolism and receptor glycosylation, while high FSS promotes adhesion and extracellular matrix interactions through integrins and laminins ^39^. Temporal analyses further demonstrated that FSS modulates transcription factors linked to oxidative stress, angiogenesis, and inflammation across different EC types ^40^. These findings align with our proteomic data, where FSS reshaped HUVEC morphology and function, shifting them from a proliferative to a structurally adaptive phenotype.

Proteomic profiling of sEVs confirmed enrichment in proteins associated with vesicle trafficking, adhesion, and structural organization, in line with previous datasets ^41^. Supporting our in vitro findings, GO enrichment analysis revealed that sEV proteins are associated with cadherin binding. Notably, several studies have reported that tumor-derived sEVs transport miRNAs targeting cadherins, thereby promoting tumor progression through epithelial–mesenchymal transition and modulation of the tumor microenvironment ^42^. Interactome analysis revealed that CME and CavME are central routes for sEVs uptake, with integrins and early endosome markers (RAB5 and EEA1) serving as critical nodes. These results suggest that internalized sEVs are rapidly targeted to the endo-lysosomal system, although FSS has been reported to redirect them away from degradation, enhancing cargo delivery and transendothelial transport ^43^.

The integrins ITGβ1 and ITGα3, present in both HUVEC and sEVs, are known to promote adhesion, vascular permeability, and metastasis ^44,45^. Interestingly, the neuroblast differentiation-associated protein AHNAK was identified in sEVs samples and exhibited potential interactions with HUVECs under both static and FSS conditions. Previously, our group reported that AHNAK expression correlates positively with the aggressiveness of breast cancer cell lines and modulates the release of EVs capable of interacting with fibroblasts, enhancing their migratory behavior ^46^. This suggests that AHNAK-containing tumor-derived sEVs may also modulate endothelial cell function, highlighting a potential cross-talk mechanism between tumor cells and the vascular endothelium that warrants further investigation. We also identified MYO6, enriched in static HUVEC, which interacts with MYL6 carried by TNBC-derived sEVs. MYL6 is linked to migration, endocytosis, and increased endothelial permeability, and has been detected in patient plasma EVs across cancer types ^47–49^. Together, these findings indicate that tumor-derived sEVs may exploit integrin signaling and MYO6-MYL6 interactions to modulate EC function, with early endosomes as key intermediates, highlighting potential biomarkers and mechanisms facilitating metastasis.

The pharmacological inhibition with MβCD and Pitstop2 significantly impaired sEVs internalization, implicating CavME and CME, respectively. Notably, siRNA-mediated knockdown of CAV-1 or CLHC reduced uptake in static cultures but paradoxically enhanced internalization under FSS, indicating that flow-dependent mechanisms can compensate for the loss of canonical endocytic pathways. Moreover, under FSS, sEVs accumulated at endothelial junctions, co-localizing with VE-CAD and interacting with CLDN-5, suggesting a dual impact on adherens and tight junctions that may weaken barrier integrity and facilitate vascular permeability.

Dynasore inhibits the GTPase activity of dynamin, thereby blocking both CME and CavME^50^. Pitstop2 interferes with binding to the N-terminal domain of clathrin, selectively inhibiting CME^51^. MβCD, in turn, removes cholesterol from the plasma membrane, blocking lipid raft–mediated endocytosis and CavME ^36^. However, these inhibitors can also affect other processes that compromise cell physiology ^52^. Without acid wash, we did not detect changes in TF-AF488 uptake by ECs, although higher doses of Dynasore and MβCD disrupted cytoskeletal organization. Most studies using HUVECs do not show cell morphology after treatment with these inhibitors; instead, they report only quantification or fluorescence micrographs with nuclear staining and the molecule of interest ^53–55^. Thus, it remains unclear whether the observed effects are due to specific drug action or to secondary effects such as membrane destabilization, cell retraction with reduced area, and cytoskeletal rearrangements.

Acid wash is a method used to remove membrane-bound molecules in endocytosis assays, thereby highlighting intracellular labeling ^56^. This strategy has been applied in HUVECs to detect VE-CAD internalization, although those data were obtained in isolated cells rather than confluent monolayers, which better reflect in vivo endothelial architecture ^57^. In our study, acid wash allowed us to select a suitable concentration of MβCD for subsequent experiments, but TF-AF488 internalization was not reduced after treatment with the other drugs. Moreover, in some fields, we observed loss of cell–cell contact. In HeLa cells, acid wash has also been reported to reduce cell–cell adhesion and viability ^58^. It is therefore possible that HUVECs behaved similarly, highlighting the limitations of this approach for studying endocytosis.

We next combined acid wash with the analysis of sEVs uptake by HUVECs. This strategy revealed that CME and CavME may contribute to sEVs internalization, since MβCD and Pitstop2 reduced vesicle uptake. The same MβCD concentration was shown to inhibit sEVs uptake from glioblastoma cells by 60% in HUVECs ^59^. While Pitstop2 has not previously been reported to inhibit sEVs uptake in HUVECs, it has been observed in pancreatic cancer, colon cancer, and trophoblast cells ^60–62^. In our hands, combining inhibitors, acid wash, and sEVs attenuated cellular damage, as evidenced by increased sEVs uptake after 4h15min of continuous Pitstop2 exposure. However, given the observed cytotoxic effects, we chose to discontinue the use of acid wash in subsequent assays.

We also compared sEVs internalization after silencing CLHC and CAV-1 in HUVECs under static and FSS conditions. In other cell types, external factors such as substrate stiffness, compression, and osmotic pressure can inhibit endocytosis ^63,64^. From a biophysical perspective, vessel topology and blood/lymph flow could be sufficient to impair CME. However, this pathway is essential for EC function. Under static conditions, HUVECs exhibit an endocytic profile similar to fibroblasts ^65^, which may explain the reduced sEVs uptake after inhibition of both CME and CavME, in contrast to cells under FSS. In a study of sEVs uptake, silencing CLHC in PC12 cells reduced sEVs internalization, while CavME inhibition had no effect ^66^. Conversely, in HeLa cells, CavME inhibition reduced sEVs uptake, whereas CME inhibition did not ^67^. Together with our findings, these studies indicate that different cell types rely on distinct endocytic routes to internalize sEVs. Specifically, both CavME and CME play crucial roles in sEVs uptake in HUVECs under static conditions.

Despite being widely used, static culture often fails to replicate the in vivo endothelial microenvironment. For this reason, we compared CLHC and CAV-1 knockdown effects also in HUVECs exposed to FSS. While knockdown reduced sEVs uptake under static conditions, it produced the opposite effect under FSS. Previous studies reported that FSS drives CAV-1 translocation to the upstream region of laminar flow ^68,69^. In contrast, HAECs exposed to 20 dyn/cm² FSS showed caveolae-rich regions polarized downstream of flow, in agreement with our HUVEC data ^70^. As noted, caveolins act as mechanosensors and plasma membrane reservoirs. Oscillatory shear stress can impair HUVEC function by reducing CAV-1, KLF2, and eNOS expression compared with physiological FSS ^71^. Consistent with this, Nawara et al. observed that 10 dyn/cm² FSS increased the formation of clathrin-coated vesicles and the rate of this process in HUVECs ^72^. Given the importance of these molecules, physiological shear may compensate for the knockdown of CAV-1 and CLHC, leading to enhanced sEVs uptake compared with controls. To date, few studies have examined HUVEC-sEVs interactions, underscoring the need to elucidate the roles of CLHC and CAV-1 in EC responses to FSS, beyond CME and CavME, and their relevance for tumor EV uptake.

Although knockdown did not alter sEVs uptake patterns in bifurcations and convergences with complex FSS dynamics, we observed spatial variation under control conditions. sEVs-HUVEC interactions were more prominent in low-FSS regions (∼4.5 dyn/cm²), whereas siCAV-1 reduced uptake in bifurcations and siCLHC abolished this variation. Interestingly, in vivo studies in zebrafish and murine models show that low FSS promotes CTC adhesion to the endothelium ^34^. This adhesion depends on integrins (e.g., αvβ3, α5β1), CD44, and laminins, whose deposition is enhanced by FSS ^73^. Moreover, tumor sEVs promote metastasis formation in the zebrafish caudal plexus, a region of low FSS ^74^. Based on these data, we hypothesize that under control conditions, sEVs preferentially interact with ECs in low-FSS regions (e.g., capillaries). By carrying integrins, as shown in our in silico data, sEVs may enhance endothelial adhesion to CTCs at these sites. In contrast, knockdown likely perturbed endocytic dynamics, altering the uptake pattern seen in controls.

Finally, we observed sEVs associated with HUVEC adherens and tight junctions. sEVs from TNBC cells are enriched in miR-939, which downregulates VE-CAD in HUVECs, thereby increasing permeability and transendothelial migration ^75^. Likewise, EVs from TNBC patients treated with doxorubicin and cyclophosphamide are enriched in phosphatidylserines and induce loss of endothelial barrier ^76^. Tumor EVs can also disrupt tight junctions; for example, colon cancer EVs increase permeability via modulation of VEGFR2, ZO-1, occludins, and CLDN5 through miR-25-3p ^77^. None of these studies, however, reported direct sEVs co-localization and interaction in endothelial junctions, as we observed. This positioning may be linked to increased permeability, ultimately facilitating transendothelial migration.

The interaction of tumor sEVs with endothelial junctions may also hold therapeutic implications. EVs derived from mesenchymal stem cells (MSCs) and brain endothelial cells have been shown to reduce blood–brain barrier (BBB) permeability in ischemic stroke models by preventing the endocytosis of ZO-1 and CLDN5 via CavME, with MSC-EVs exerting a stronger protective effect ^78^. Similarly, Shao et al. (2025) demonstrated that MSC-derived EVs alleviate sepsis by limiting neutrophil extracellular trap (NET) formation and maintaining adherens junction integrity in lung ECs through calpain-1/2 regulation ^79^. Given that NETs contribute to metastasis and tumor EVs have been implicated in NET induction ^80^, engineering EVs to counteract NET formation could represent a novel strategy to prevent metastatic dissemination while preserving endothelial barrier function.

Despite the insights gained, this study has some limitations. First, the in vitro models, while incorporating FSS and better recapitulating the in vivo behavior of ECs, cannot fully recapitulate the complex hemodynamic and biochemical environment of the vasculature in vivo, including interactions with perivascular cells and the extracellular matrix. Second, pharmacological inhibitors and siRNA-mediated knockdowns, while informative, may have off-target effects or trigger compensatory mechanisms, particularly under FSS conditions, complicating interpretation of endocytic pathway contributions. Finally, the study primarily focused on short-term uptake and junctional interactions, leaving long-term functional consequences on endothelial barrier integrity and tumor cell transmigration unaddressed. Future studies employing in vivo models and more refined EV subpopulation isolation will be essential to validate and extend these findings.

Despite these limitations, our study provides important mechanistic insights into the interaction between TNBC-derived sEVs and endothelial cells, emphasizing the pivotal roles of CME, CavME and the preferential targeting of adherens and tight junctions under fluid shear stress. By integrating in silico and in vitro analyses, we show that CME and CavME mediate sEV uptake under both static and FSS conditions, offering a mechanistic framework that could guide nanoparticle-based therapeutic strategies, harnessing tumor EV markers and endocytic pathways for the precise delivery of nucleic acids, proteins, or drugs. Collectively, these findings lay the groundwork for future studies exploring EV-mediated vascular remodeling, metastatic dissemination, and the development of EV-based therapeutic interventions.

## 4. Methods

### 4.1. Reagents

Most chemical reagents were purchased from Sigma-Aldrich. Reagents obtained from other suppliers are specified individually. Antibody details, including catalog number, application, and working concentration, are listed in Table 1.

**Table 1.**
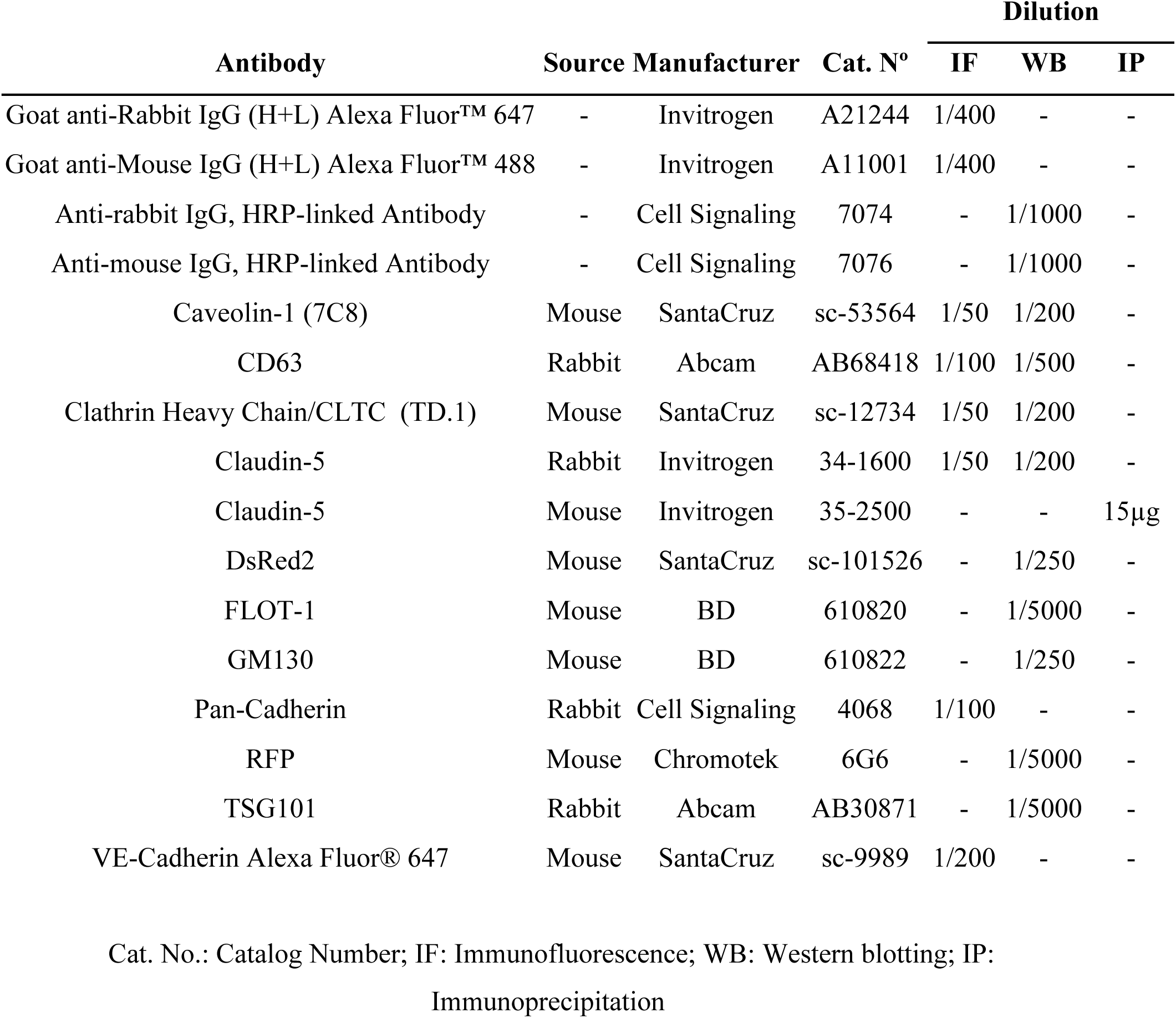
List of antibodies used in this study.

| Antibody | Source | Manufacturer | Cat. N° | Dilution |  |  |
| --- | --- | --- | --- | --- | --- | --- |
|  |  |  |  | IF | WB | IP |
| Goat anti-Rabbit IgG (H+L) Alexa Fluor™ 647 | - | Invitrogen | A21244 | 1/400 | - | - |
| Goat anti-Mouse IgG (H+L) Alexa Fluor™ 488 | - | Invitrogen | A11001 | 1/400 | - | - |
| Anti-rabbit IgG, HRP-linked Antibody | - | Cell Signaling | 7074 | - | 1/1000 | - |
| Anti-mouse IgG, HRP-linked Antibody | - | Cell Signaling | 7076 | - | 1/1000 | - |
| Caveolin-1 (7C8) | Mouse | SantaCruz | sc-53564 | 1/50 | 1/200 | - |
| CD63 | Rabbit | Abcam | AB68418 | 1/100 | 1/500 | - |
| Clathrin Heavy Chain/CLTC (TD.1) | Mouse | SantaCruz | sc-12734 | 1/50 | 1/200 | - |
| Claudin-5 | Rabbit | Invitrogen | 34-1600 | 1/50 | 1/200 | - |
| Claudin-5 | Mouse | Invitrogen | 35-2500 | - | - | 15µg |
| DsRed2 | Mouse | SantaCruz | sc-101526 | - | 1/250 | - |
| FLOT-1 | Mouse | BD | 610820 | - | 1/5000 | - |
| GM130 | Mouse | BD | 610822 | - | 1/250 | - |
| Pan-Cadherin | Rabbit | Cell Signaling | 4068 | 1/100 | - | - |
| RFP | Mouse | Chromotek | 6G6 | - | 1/5000 | - |
| TSG101 | Rabbit | Abcam | AB30871 | - | 1/5000 | - |
| VE-Cadherin Alexa Fluor® 647 | Mouse | SantaCruz | sc-9989 | 1/200 | - | - |

Cat. No.: Catalog Number; IF: Immunofluorescence; WB: Western blotting; IP: Immunoprecipitation

### 4.2. Cell culture

The MDA-MB-231 cell line was provided by the American Type Culture Collection (ATCC) and it was cultivated with DMEM (Sigma Chemical Co.) supplemented with 5% fetal bovine serum (FBS; Cultilab) and 1% penicillin/streptomycin (Sigma Chemical Co.). HUVEC (Lonza) were cultivated with EGM-2 Endothelial Cell Growth Basal Medium supplemented with its specific bullet kit (Lonza). Cell flasks were kept in a humidified incubator at 37°C and 5% CO2. Medium exchanges were performed every two days.

### 4.3. Cell aligning using orbital shaker

HUVECs were seeded at a density of 50,000 cells/cm² in 100 mm Petri dishes. Upon reaching confluence, the medium was replaced with culture medium supplemented with 4% dextran (molecular weight: 450,000–650,000 g/mol). Cultures were then subjected to FSS using an orbital shaker (Thermolyne Rotomix 50800 Orbital Shaker) placed inside a humidified incubator at 37°C and 5% CO₂. Control HUVECs were cultured in parallel at the same density without FSS induction.

The wall shear stress (τmax, dyn/cm²) was calculated using the equation:

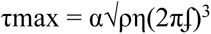

where α is the orbital radius (1 cm), ρ is the medium density (∼1 g/mL), η is the viscosity of the 4% dextran-supplemented medium (0.06 poise), and f is the rotational frequency (s⁻¹) ^81,82^.

Based on this equation, an FSS of approximately 20 dyn/cm² was achieved at a rotation speed of 173.4 rpm, which corresponds to physiological levels of macrocirculatory shear stress. After 96 h of incubation, the morphology of HUVECs in the periphery of the petri dish was assessed by phase-contrast microscopy, then, aligned cells were lysed. Cell circularity was measured as the ratio between cell length and width using FIJI/ImageJ software (NIH, version 1.54r). Values close to 1.0 indicate higher circularity, whereas values approaching 0 reflect a flow aligned morphology.

### 4.4. Trypsin Digestion in Solution

Protein samples (100 μg from cell lysates and 3μg from EV lysates) were denatured with 6 M guanidine hydrochloride (GuHCl) to a final concentration of 3 M, followed by addition of dithiothreitol (DTT, 5 mM final). Samples were incubated at 37 °C for 1h and then alkylated with iodoacetamide (IAA, 15 mM final) for 30min at room temperature. Proteins were precipitated with cold acetone (8 volumes) and methanol (1 volume) at −80°C for 2 h. After centrifugation (14,000 × g, 10 min), protein pellets were washed twice with cold methanol and resuspended in 10 mM NaOH. Samples were then diluted with HEPES buffer (50 mM, pH 7.5) to a final volume of 100 μL. Trypsin was added at an enzyme-to-substrate ratio of 1:100, and digestion was carried out at 37 °C for 18 h. Tryptic peptides were desalted using C18 StageTips, dried in a SpeedVac, and stored at −80°C until LC–MS/MS analysis. Prior to injection, samples were reconstituted in 50 μL of 0.1% formic acid.

### 4.5. Mass Spectrometry (LC–MS/MS)

Peptide separation was performed using an Easy-nLC 1200 UHPLC system (Thermo Scientific) with a binary gradient of solvent A (0.1% formic acid) and solvent B (80% acetonitrile, 0.1% formic acid). Peptides were first loaded onto a trap column (Acclaim PepMap 100, C18, 3 μm, 75 μm × 2 cm, Thermo Scientific) with 20 μL of solvent A at 500 bar. Elution was carried out on an analytical column (Acclaim PepMap RSLC, C18, 2 μm, 75 μm × 15 cm, Thermo Scientific) at 300 nL/min using the following gradient: 5–28% solvent B over 80 min, 28–40% B over 10 min, ramp to 90% B in 2 min, followed by a 12 min wash. The system was re-equilibrated with solvent A before each injection.

Mass spectrometry was performed on an Orbitrap Fusion Lumos (Thermo Scientific) equipped with a Flex NG nanospray ion source, operated in positive ESI mode (capillary temperature 300°C, S-Lens RF level 30%). Full MS scans were acquired in the Orbitrap at a resolution of 120,000 (m/z 200), AGC target 5 × 10⁵, scan range m/z 350–1550. Data-dependent MS2 scans were acquired in 3 s cycles with Orbitrap resolution of 30,000 (m/z 200), maximum injection time 54ms, isolation window 1.2 m/z, and dynamic exclusion of 40 s. Higher-energy collisional dissociation (HCD) was performed at 30% normalized collision energy. The APD algorithm was enabled to improve precursor detection. Mass calibration was performed before each run using the EASY-IC™ function.

### 4.6. Proteomics Data Processing

Raw data files were processed with MaxQuant v2.0.3.0 using a 1% false discovery rate (FDR) for both protein and peptide identifications. Spectra were searched against the UniProt/SwissProt Homo sapiens database (release 2022_10; 42,414 entries), supplemented with 245 common contaminants and reversed decoy sequences. Trypsin specificity was set with up to two missed cleavages. Carbamidomethylation of cysteine was defined as a fixed modification, while methionine oxidation, asparagine/glutamine deamidation, and N-terminal acetylation were included as variable modifications. Precursor mass tolerance was set at 4.5ppm and fragment mass tolerance at 20 ppm.

Label-free quantification (LFQ) was performed using the MaxLFQ algorithm, with “match between runs” and “re-quantify” enabled. Protein intensities from proteinGroups.txt were log₂-transformed and quantile-normalized using the ‘preProcessCore’ package in R. Differential expression between HUVEC Static and HUVEC FSS conditions was assessed using Student’s t-test. Proteins with p ≤ 0.05 and |log₂ fold change| > 1.5 were considered differentially abundant. Reproducibility among biological replicates was evaluated by Pearson correlation (data not shown).

Gene Ontology (GO) enrichment analysis of differentially expressed proteins was performed using the pipeline described by Bonnot et al. (2019), and visualizations were generated with ggplot2 in R. The same approach was applied to the sEVs dataset ^83^.

### 4.7. Protein-Protein Interactions

Differentially expressed HUVEC proteins were annotated for enrichment in the GO term Endocytosis (GO:0006897). For EVs, identified proteins were cross-referenced with the Vesiclepedia database, including the top 100 most detected proteins in EVs ^84^. Venn diagrams were generated using FunRich ^85–87^.

Protein–protein interaction networks were built with Cytoscape v3.8.2 ^88^. using protein lists from HUVEC Static, HUVEC FSS and sEVs. Interactions were retrieved from the STRING database (physical interactors mode, cut-off 0.7, unrestricted number of interactions) ^89^. Networks were merged, and hub proteins were identified using the CytoHubba plugin, ranking the top 10 nodes by maximal clique centrality (MCC) ^90^.

### 4.8. Transformation of E. coli DH5-alpha and Transfection of MDA-MB-231 Cell Line with Plasmid DNA

The plasmids pKT2/CAGXSP/CD63mScarlet (Plasmid #182972, Addgene) and pCMV(CAT)T7-SB100 (Plasmid #34879, Addgene) were purchased from Addgene ^91,92^. Competent *E. coli* DH5-alpha cells (Sigma-Aldrich) were thawed on ice and incubated with 100 ng/µL of purified plasmid DNA for 30min. Cells were heat-shocked at 42°C for 42s, cooled on ice for 2 min, and recovered in 1 mL LB medium without antibiotics at 37°C for 1h under agitation. Cultures were centrifuged at 800 × g for 1 min at room temperature, and the pellet was resuspended in ∼100µL of LB. Aliquots were plated on LB agar (2%) containing 50 µg/mL kanamycin (for pKT2/CAGXSP/CD63mScarlet) or 25 µg/mL chloramphenicol (for pCMV(CAT)T7-SB100), followed by incubation for 16h at 37°C. Single colonies were grown in LB medium supplemented with the respective antibiotics, and glycerol stocks (1:1 with 50% glycerol) were stored at –80°C. Plasmid DNA was isolated using the PureYield™ Plasmid Midiprep System (Promega, Cat. A2495) and quantified with the Epoch Microplate Spectrophotometer (Agilent Technologies).

MDA-MB-231 cells were seeded in 6-well plates at a density of 10,500 cells/cm² and cultured overnight at 37°C in a 5% CO₂ atmosphere. Cells were transfected with the isolated plasmids using Lipofectamine 3000 (ThermoFisher, Cat. L3000001) in Opti-MEM™ Reduced Serum Medium (ThermoFisher, Cat. 31985070), following the manufacturer’s instructions. A plasmid ratio of 2 µg pKT2/CAGXSP/CD63mScarlet to 1 µg pCMV(CAT)T7-SB100 was used per reaction. After 5 h, the medium was replaced with DMEM supplemented with 10% FBS, and cells were cultured until 90% confluence. Stable expression of CD63-mScarlet was achieved by selection with 4 µg/mL puromycin. The culture was expanded from cell clones, cryopreserved, and sorted to isolate a population with high CD63-mScarlet expression using a BD FACSDiscover S8 cell sorter (BD Biosciences).

For validation of fluorescence, MDA-MB-231 WT and MDA-MB-231 CD63-mScarlet cells were cultured as previously described, trypsinized, and transferred to 12 × 75 mm polypropylene tubes (BD Falcon). Cells were then centrifuged at 1500 rpm for 5 minutes at 4 °C and resuspended in 200 µL of buffer (PBS containing 2% FBS). Samples were immediately acquired on a Cytek Aurora flow cytometer (Cytek Biosciences).

Data of 100,000 events/condition were analyzed using FlowJo software, version 7.5.5 (Tree Star), according to the gating strategy illustrated in Supplementary Figure 1. Briefly, small-sized and low-complexity events (FSC and SSC, respectively) were excluded as cellular debris (Supplementary Figure 1A). Doublets were removed based on forward scatter area (FSC-A) and height (FSC-H) parameters (Supplementary Figure 1B). Subsequently, within the single-cell population, CD63-mScarlet–positive cells were identified (Supplementary Figure 1C).

### 4.9. EVs isolation and characterization

MDA-MB-231 cells stably expressing CD63 fused with mScarlet were seeded at a density of 27,000 cells/cm² and maintained at 37°C in a 5% CO₂ atmosphere for 24 h. The following day, cells were washed twice with PBS and incubated in Opti-MEM™ Reduced Serum Medium (ThermoFisher) for 48 h. Conditioned medium was collected and sequentially centrifuged at 300 × g for 10 min, 2,000 × g for 25 min, and 10,000 × g for 30 min at 4°C using a 5804 R refrigerated centrifuge (Eppendorf) to remove suspended cells, cellular debris, and large EVs, respectively.

The supernatant was concentrated up to 30 mL by Centricon Plus Centrifugal Filter (MWCO 100 kDa, Sigma Aldrich) and transferred into UltraClear 25 × 89 mm tubes (Beckman Coulter, Cat. 344058), and 2 mL of 60% iodixanol was carefully underlaid using a 5mL syringe fitted with a long metal needle. Tubes were ultracentrifuged at 100,000 × g for 4 h at 4°C in an SW28 rotor (Optima XPN-100 ultracentrifuge, Beckman Coulter), resulting in particle enrichment near the 60% iodixanol interface. A 3 mL fraction (2 mL 60% iodixanol + 1 mL culture medium containing EVs) was collected from the bottom and homogenized to yield a 40% iodixanol solution. Discontinuous iodixanol gradients were prepared by layering 10%, 20%, and 40% iodixanol solutions in sucrose buffer (0.25 M sucrose, 10 mM Tris-base, pH 7.5) into 14 mL, 14 × 95mm tubes (Beckman Coulter, Cat. 344060), followed by 5% iodixanol on top, reaching a final volume of 12mL. Gradients were ultracentrifuged at 100,000 × g for 20 h at 4°C in an SW41 rotor (L8-80 ultracentrifuge, Beckman Coulter). Columns were fractionated into twelve 1 mL fractions, transferred into 3.5 mL tubes (13 × 51mm, Beckman Coulter, Cat. 349622), resuspended in 2 mL of ice-cold PBS, and ultracentrifuged again at 100,000 × g for 2 h at 4°C in a TLA110 rotor (Optima MAX tabletop ultracentrifuge, Beckman Coulter).

For molecular validation, aliquots corresponding to 10⁵ particles from Fraction 6 were collected from each resuspended fraction. Whole-cell lysates (WCL) of MDA-MB-231 CD63-mScarlet cells, cultured under the same conditions, were prepared in RIPA buffer and quantified with the Pierce™ BCA assay (ThermoFisher, Cat. A550864). A total of 5 µg of WCL protein was used for Western blotting. Membranes were probed with antibodies against positive EV markers (TSG101, FLOT-1, CD63), a negative marker (GM130), and anti-Red Fluorescent Protein (RFP) to confirm the presence of mScarlet, derived from RFP ^93^. EV size and concentration were determined by nanoparticle tracking analysis (NTA) using the ZetaView® x30 instrument (Particle Metrix). EVs obtained through this protocol were designated as sEVs.

### 4.10. Adhesion of sEVs to Glass Coverslips

Glass coverslips (13 mm) were coated overnight at 4°C with 100 µg/mL Poly-D-Lysine Hydrobromide (PDL). After washing three times with PBS, 10⁸ sEVs particles in 50 µL PBS were added per coverslip and incubated overnight at 4°C, protected from light. Coverslips were fixed with 4% paraformaldehyde (PFA) in PBS for 10 min, immunostained with CD63 and goat anti-rabbit AlexaFluor 488, mounted, and imaged using a Nikon A1R confocal microscope equipped with a CFI Apochromat TIRF 60X Oil.

### 4.11. Immunogold Labeling of sEVs for Transmission Electron Microscopy

A total of 2 × 10⁸ sEVs suspended in 20 µL of PBS were adsorbed onto nickel grids coated with formvar and carbon (Koch Electron Microscopy LTDA, FCF200-400 MESH-NI-50) for 1 h at room temperature. Excess liquid was gently removed using filter paper, and the grids were blocked with 10% donkey serum in PBS for 10min. Subsequently, the grids were washed twice for 2 min each in droplets of 1% bovine serum albumin (BSA) prepared in filtered PBS (0.22 µm). For immunolabeling, grids were incubated overnight at room temperature in a humid chamber with 20 µL of anti-CD63 antibody diluted in PBS containing 1% BSA. After incubation, grids were washed eight times for 2 min each in PBS and then incubated for 2 h at room temperature in a humid chamber with donkey anti-mouse or anti-rabbit IgG conjugated to 18 nm gold particles (Jackson ImmunoResearch, 715-215-150) diluted 1:10 in PBS containing 1% BSA. Following secondary labeling, grids were washed eight times for 2 min each in PBS, fixed with 2.5% glutaraldehyde for 10 min, and rinsed three times for 2 min in PBS. The grids were then air-dried for 1 h at room temperature and counterstained with 2% uranyl acetate for 5 min. Micrographs were acquired using a JEOL 1010 transmission electron microscope.

### 4.12. Western Blot

Protein samples were denatured at 95°C in loading buffer (0.5 mM Tris-HCl pH 6.8, 10% SDS, 10% glycerol, 715 mM β-mercaptoethanol, 0.05% bromophenol blue) and resolved by SDS-PAGE. Gels consisted of a resolving layer (bis-acrylamide concentration adjusted to the protein(s) of interest, 375 mM Tris-HCl pH 8.8, 0.1% SDS) and a 3% stacking layer (125 mM Tris-HCl pH 6.8, 0.1% SDS). Electrophoresis was performed at 100 V for 1.5 h in running buffer (25 mM Tris-HCl pH 8.3, 200 mM glycine, 0.1% SDS).

Proteins were transferred to nitrocellulose membranes using transfer buffer (25 mM Tris, 193 mM glycine, 20% methanol) at 18 V for 16 h. Membranes were blocked in 5% nonfat dry milk in TBS (10 mM Tris-HCl pH 7.4, 0.9% NaCl) for 1 h at room temperature, then incubated with primary antibodies (monoclonal or polyclonal, depending on the experiment). After three washes in TBS-T (10mM Tris-HCl pH 7.4, 0.9% NaCl, 0.2% Tween-20), membranes were incubated for 1 h at room temperature with HRP-conjugated goat anti-rabbit or goat anti-mouse secondary antibodies, followed by three additional washes in TBS-T.

Detection was performed with Clarity™ Western ECL substrate (Bio-Rad, Cat. 1705060) and imaged using a ChemiDoc system (Bio-Rad). Band intensities were quantified with FIJI/ImageJ software (NIH, version 1.54r).

### 4.13. Crystal violet staining

HUVECs were seeded at a density of 15,000 cells/cm² in 96-well plates and maintained at 37°C in a humidified atmosphere containing 5% CO₂ until reaching confluence. Cells were then treated for 4 h with Dynasore (100, 50, or 25 µM; Sigma-Aldrich, Cat. D7693), Pitstop2 (30, 20, or 10 µM; Abcam, Cat. Ab120687), or methyl-β-cyclodextrin (MβCD; 1.25, 2.5, or 5 µg/mL; Sigma-Aldrich, Cat. 332615). As controls, DMSO was used at concentrations equivalent to those in the highest doses of Dynasore (0.2%) and Pitstop2 (0.1%).

After treatment, monolayers were washed with PBS and incubated with 50 µL of 0.5% crystal violet for 20min at room temperature. Wells were rinsed briefly with ddH₂O, and plates were air-dried overnight at room temperature. The stain was solubilized with 200 µL methanol, and absorbance was measured at 570 nm using a microplate reader.

### 4.14. Transferrin Endocytosis Assay with Endocytosis Inhibitors

HUVECs were seeded at a density of 50,000 cells/cm² on glass coverslips in 24-well plates and maintained at 37°C in a humidified atmosphere containing 5% CO₂ until reaching confluence. Cells were then treated for 30 min with 100 μM Dynasore and 2.5 μg/mL MβCD, or 15min with 30 μM Pitstop2. As controls, cells were treated with DMSO at concentrations corresponding to the highest levels used for Dynasore (0.2%) and Pitstop2 (0.1%).

Following this incubation period, SEVs isolated from the MDA-MB-231 cell line expressing the CD63-mScarlet fusion protein were added at a concentration of 5 × 10⁸ particles/mL, diluted in culture medium containing the respective endocytosis inhibitor, and incubated for 30 min. After incubation, cells were subjected to an acid wash using 0.1 M glycine buffer (pH 2.2), followed by washing with culture medium. Cells were then fixed with 4% paraformaldehyde (PFA), permeabilized with 0.5% Triton X-100 in PBS, and stained for the cytoskeleton using phalloidin conjugated to Alexa Fluor™ 488 (Cat. A12379, Invitrogen). Nuclei were stained with Hoechst. Coverslips were mounted and analyzed using a Nikon A1R confocal microscope. Three PBS washes were performed between each step, followed by a final wash with ddH₂O. TF-AF488 uptake was quantified using FIJI/ImageJ software (NIH, version 1.54r).

### 4.15. sEVs Endocytosis Assay with siRNA Knockdown

HUVECs were trypsinized, counted, and 7.5 × 10⁵ cells were transfected with 100 nM small interfering RNA (siRNA) targeting CLHC or CAV-1, or with a scrambled sequence as a negative control (Santa Cruz Biotechnology). Transfection was performed according to the manufacturer’s instructions using the HUVEC Nucleofector™ Kit (Lonza, Cat. VPB-1002) and the Nucleofector® 2b Device (Lonza). Transfection efficiency was confirmed in parallel by Western blotting, as previously described, using monoclonal anti-CLHC and anti-CAV-1 antibodies (Table 1).

Following transfection, 2.0 × 10⁵ cells were seeded in µ-Slide y-shaped channels (FSS condition; Ibidi, Cat. 80126) and 1.0 × 10⁵ cells were seeded on glass coverslips in 24-well plates (static condition). Cells were incubated for 4 h at 37°C in a humidified atmosphere containing 5% CO₂, followed by medium replacement. For the static condition, the medium was replaced again after 24 h. For the FSS condition, channels were connected to the Ibidi Pump System (Ibidi, Cat. 10902) with 13 mL of culture medium 24 h after transfection and exposed to laminar flow at 8.5 dyn/cm² for 1 h to allow adaptation. The FSS was then increased to 20 dyn/cm² for 23 h and subsequently reduced to 8.5 dyn/cm². At this stage, 5 × 10⁸ particles/mL of sEVs isolated from MDA-MB-231 CD63-mScarlet cells were added, followed by 4h incubation in both static and FSS conditions.

Cells were fixed with 4% PFA, then permeabilized and blocked with PBS containing 3% goat serum, 0.3% Triton X-100, and 0.05% Tween 20. Samples were incubated overnight at 4 °C in a humidified chamber with anti-PAN-CAD antibody, followed by staining with Alexa Fluor™ 647-conjugated goat anti-rabbit secondary antibody and DAPI for nuclei. Coverslips and slides were mounted and imaged using a ZEISS Axio Observer 7 epifluorescence microscope equipped with Plan-Apochromat 63x/1.40 Oil objective (Zeiss) or Leica Stellaris WLL equipped with HC PL APO 63x/1.40 OIL CS2 objective (Leica).

sEVs internalization was quantified using CellProfiler 4.2.8 (Broad Institute) with the CellPose algorithm ^94–98^.

### 4.16. Co-immunoprecipitation

HUVEC were seeded and exposed to FSS using an orbital shaker, as previously described, while control cells were maintained under static conditions. Cells were then incubated for 4 h with 5 × 10⁸ particles/mL of sEVs isolated from MDA-MB-231 CD63-mScarlet cells. After incubation, cells were lysed in 1% NP-40 prepared in TBS, and protein concentration was determined using the BCA assay. Co-immunoprecipitation (Co-IP) was performed with the Dynabeads kit (Thermo Fisher) according to the manufacturer’s protocol, using 15 μg of either anti-CLDN-5 or anti-Pan-CAD antibody. Samples corresponding to sEVs, whole cell lysate (input), supernatant of the first wash (wash), Co-IP CLDN-5, and Co-IP Pan-CAD were separated by SDS-PAGE, transferred to nitrocellulose membranes, and probed with anti-dsRed2 and anti-CLDN-5 antibodies.

### 4.17. Statistical analysis

Normality was assessed with Kolmogorov-Smirnov or Shapiro-Wilk tests. Nonparametric comparisons used Mann-Whitney or Wilcoxon tests; parametric comparisons used Student’s t-test (paired or unpaired). Data analysis and plotting were performed in GraphPad Prism 10.0.

### 4.18. Artificial Intelligence Generated Content

ChatGPT-5.0 was used exclusively to refine and improve the clarity and readability of the manuscript text. All AI-assisted edits were carefully reviewed, verified, and approved by all co-authors prior to submission.

## Author contributions

NJV conceptualized the study, performed most of the experiments, conducted formal analyses, and wrote the initial draft of the manuscript. BHS provided training and supervised the experiments conducted at Vanderbilt University. ASY and RHGT contributed to the characterization of extracellular vesicles and provided critical input during scientific discussions. LES participated in the design and interpretation of the in silico analyses. AZ and MS performed proteomic analyses and supported the in silico work. VSB carried out the cloning of plasmids. GRT and MCC provided technical support for electron and confocal microscopy. RGJ and AW secured financial support, with AW also contributing to experimental design and data analysis. VMF supervised all experimental work, acquired funding, and critically reviewed and edited the manuscript. All authors read, revised, and approved the final version of the manuscript.

## Acknowledgements

This work was supported by the INCT 3D Model coordinated by Marimelia Aparecida Porcionatto (CAPES, Grant No. 58/2022). We thank Professors Paolo Di Mascio and Graziella E. Ronsein for the Mass spectrometry analyses performed at the Redox Proteomics Core of the Mass Spectrometry Resource at Chemistry Institute, University of Sao Paulo (FAPESP 2012/12663-1, 2016/00696-3, CEPID Redoxoma 2013/07937-8). Flow cytometry analyses were conducted at the Flow Cytometry Facility of the Department of Immunology, Institute of Biomedical Sciences, University of São Paulo (ICB/USP).Confocal imaging using the Leica Stellaris system was performed at the Core Facility for Scientific Research, University of São Paulo (CEFAP/USP).

